# The Cif proteins from *Wolbachia* prophage WO modify sperm genome integrity to establish cytoplasmic incompatibility

**DOI:** 10.1101/2022.01.15.476471

**Authors:** Rupinder Kaur, Brittany A. Leigh, Isabella T. Ritchie, Seth R. Bordenstein

## Abstract

Inherited microorganisms can selfishly manipulate host reproduction to drive through populations. In *Drosophila melanogaster*, germline expression of the native *Wolbachia* prophage WO proteins CifA and CifB cause cytoplasmic incompatibility (CI) in which embryos from infected males and uninfected females suffer catastrophic mitotic defects and lethality; however, in infected females, CifA expression rescues the embryonic lethality and thus imparts a fitness advantage to the maternally-transmitted *Wolbachia*. Despite widespread relevance to sex determination, evolution, and vector control, the mechanisms underlying when and how CI impairs male reproduction remain unknown and a topic of debate. Here we use cytochemical, microscopic, and transgenic assays in *D. melanogaster* to demonstrate that CifA and CifB proteins of *w*Mel localize to nuclear DNA throughout the process of spermatogenesis. Cif proteins cause abnormal histone retention in elongating spermatids and protamine deficiency in mature sperms that travel to the female reproductive tract with Cif proteins. Notably, protamine gene knockouts enhance wild type CI. In ovaries, CifA localizes to germ cell nuclei and cytoplasm of early-stage egg chambers, however Cifs are absent in late-stage oocytes and subsequently in fertilized embryos. Finally, CI and rescue are contingent upon a newly annotated CifA bipartite nuclear localization sequence. Together, our results strongly support the Host Modification model of CI in which Cifs initially modify the paternal and maternal gametes to bestow CI-defining embryonic lethality and rescue.

## Introduction

Numerous animal species harbor heritable microorganisms that alter host fitness in beneficial and harmful ways. The most common, maternally-inherited bacteria are *Wolbachia* that typically reside intracellularly in reproductive tissues of both male and female arthropods. Here, they induce reproductive modifications with sex specific effects such as cytoplasmic incompatibility (CI) that can selfishly drive the bacteria to high frequencies in host populations. CI also notably yields important consequences on arthropod speciation [1–3] and vector control strategies [4–9] by causing lethality of embryos from *Wolbachia*-infected males and uninfected females. As CI is rescued by *Wolbachia*-infected females with the same strain [10, 11], the phenotype accordingly imparts a relative fitness advantage to infected females that transmit the bacteria [12].

Two genes, *cytoplasmic incompatibility factors cifA* and *cifB*, occur in *Wolbachia* prophage WO within the eukaryotic association module enriched for arthropod functions and homology [13–15]. We previously demonstrated that dual, transgenic expression of *cifA* and *cifB* from *w*Mel *Wolbachia* in *Drosophila melanogaster* males induces CI, while single expression of *cifA* in females rescues CI [16, 17]. These results form the basis of the Two-by-One genetic model of CI for several, but not all, strains of *Wolbachia* [13,15,18,19]. At the cellular level, CI-defining lethality associates with chromatin defects and mitotic arrest within the first few hours of embryonic development. Normally after fertilization, the sperm-bound ‘protamines’ are removed in the embryo and replaced by maternally supplied ‘histones’, resulting in the rapid remodeling of the paternal chromatin [20]. However, during CI, there is a delay in the deposition of maternal histones onto the paternal chromatin, resulting in altered DNA replication, failed chromosome condensation, and various mitotic defects that generate embryonic death [13,21– 27].

The incipient, pre-fertilization events in the reproductive tissues that establish CI and rescue remain enigmatic and under recent debate, namely whether (i) Cifs modify the paternal genome during spermatogenesis (Host-Modification model) or embryogenesis (Toxin-Antidote model) and (ii) rescue occurs or does not occur by CifA binding CifB in the embryo [28, 29]. These two key observations can conclusively differentiate the mechanistic models of CI, though they have not been explicitly addressed to date. Notably, paternal transmission of the proteins from sperm to embryo may occur under either mechanistic model [11, 29]. Recent work proposed the Toxin-Antidote model is operational using transgenic, heterologous expression of Cif proteins from *w*Pip *Wolbachia* [30]. In this study, the CifB*_w_*_Pip_ protein paternally transfers to the fly embryo and associates with DNA replication stress of the paternal genome in the embryo. However, without the ability to rescue this non-native, transgenic CI and thus visualize CifA- CifB binding in the rescue embryo, these interesting results do not yet resolve the predictions of the two models. Additionally, the paternal DNA replication defects observed in the embryos may be established before fertilization by the Cif proteins, in concordance with Host Modification model.

Here, we develop antibodies to localize Cif proteins from *w*Mel *Wolbachia* during *D. melanogaster* gametogenesis and embryogenesis and then perform genome integrity measurements of developing sperm across transgenic, mutant, and wild type treatment groups. We describe the following cell biological and gametic chromatin events underpinning the Host Modification model of CI and rescue: (i) CifA and CifB proteins localize to the developing sperm nuclei from early spermatogonium stage to late elongating spermatids; (ii) In mature sperm, CifA associates with sperm tails and occasionally occurs in the acrosome, whereas CifB localizes to the acrosome in all mature sperms; (iii) Cifs increase histone retention in developing spermatids and decrease protamine levels in mature sperms; (iv) Both CI and rescue are dependent upon a newly annotated bipartite nuclear localization signal (bNLS) in CifA that impacts nuclear localization and sperm protamine levels; (v) During copulation, both Cif proteins transfer with the mature sperm exhibiting reduced protamine levels; importantly, protamine mutant flies enhance wild type CI; (vi) In the ovaries, CifA is cytonuclear in germline stem cells and colocalizes with *Wolbachia* in the nurse cell cytoplasm; (vi) CifA is absent in the embryos, and thus rescue must be established in oogenesis independently of CifA’s presence in the embryo. Taken together, results demonstrate that prophage WO-encoded Cif proteins from *Wolbachia* invade gametic nuclei to modify chromatin integrity at the histone to protamine transition stage in males. At the mechanistic level, results support the Host Modification model of CI and rescue whereby native Cifs wield their impacts prior to fertilization.

## Results

### CifA and CifB invade sperm nuclei during spermatogenesis and spermiogenesis

To evaluate the cellular localization of the Cif proteins, we generated monospecific polyclonal antibodies for visualizing the proteins in reproductive tissues (Fig S1). In *D. melanogaster* males, the sperm morphogenesis process is subdivided into two events (i) Spermatogenesis including mitotic amplification and meiotic phases, and (ii) Spermiogenesis, a post-meiotic phase. During spermatogenesis, the germline stem cell undergoes four rounds of synchronous mitotic divisions to produce 16 precursor cells called spermatogonia. The spermatogonia then grow and become spermatocytes [31]. After the growth phase, the spermatocytes divide by meiosis and differentiate into 64 haploid round onion spermatids. Post-meiosis, the round sperm nuclei elongate to gradually change their shape accompanied by reorganization of the chromatin during the canoe stage [32]. This results in an individualization complex forming slim, needle-shaped sperm nuclei with reduced volume [31, 32]. Elongation and individualization of the spermatids is the final stage of spermiogenesis, after which the mature sperms are transported to the seminal vesicle [31, 33].

CifA, but not CifB, localizes in the germline stem cells at the apical end of testes in <8 hr old *cifA* and *cifB* transgene-expressing (Fig 1) and wild type *w*Mel+ males (Fig S2). CifA and CifB were both detected in the nuclei of mitotic spermatogonium and spermatocytes. CifA is more abundant than CifB in the spermatogonium stage (Fig S3A, Table S1). In the post-meiotic, round onion spermatids, clusters of both CifA and CifB are adjacent to the nuclei. In the elongating canoe-shaped spermatids, CifA and CifB localize apical to the sperm head nucleus, in what is likely the acrosome (Fig 1, Fig S2). CifB is present in all of the spermatid nuclei, whereas CifA is present on average in 39% of the elongating spermatids per sperm bundle (Fig S3B). During the elongating canoe stage, chromatin-bound histones are typically removed and replaced with protamines to yield compact nuclear packaging and chromatin organization of sperm DNA [32]. After nuclear compaction is complete, neither of the Cif proteins are detectable in late spermatid needle-shaped nuclei (Fig 1) and in the mature sperms from the seminal vesicle (Fig S4), indicating either the Cif proteins are fully stripped, or they might not be accessible by the antibodies when the chromatin is tightly compacted [34, 35]. To evaluate Cif presence/absence in the mature sperm, we decondensed sperms after isolations from seminal vesicles of <8 hr old males (see methods) and stained them with the respective Cif antibodies. CifA is common along sperm tails in a speckled pattern (Fig 1, Fig S2) and infrequently present in the acrosome region, on average in 45% or 0% of the mature sperm heads depending upon the sampled seminal vesicles (Fig S3B). CifB is present in all of the acrosomal tips of the sperms and not localized to the sperm tails (Fig 1, Fig S2).

**Fig 1.**
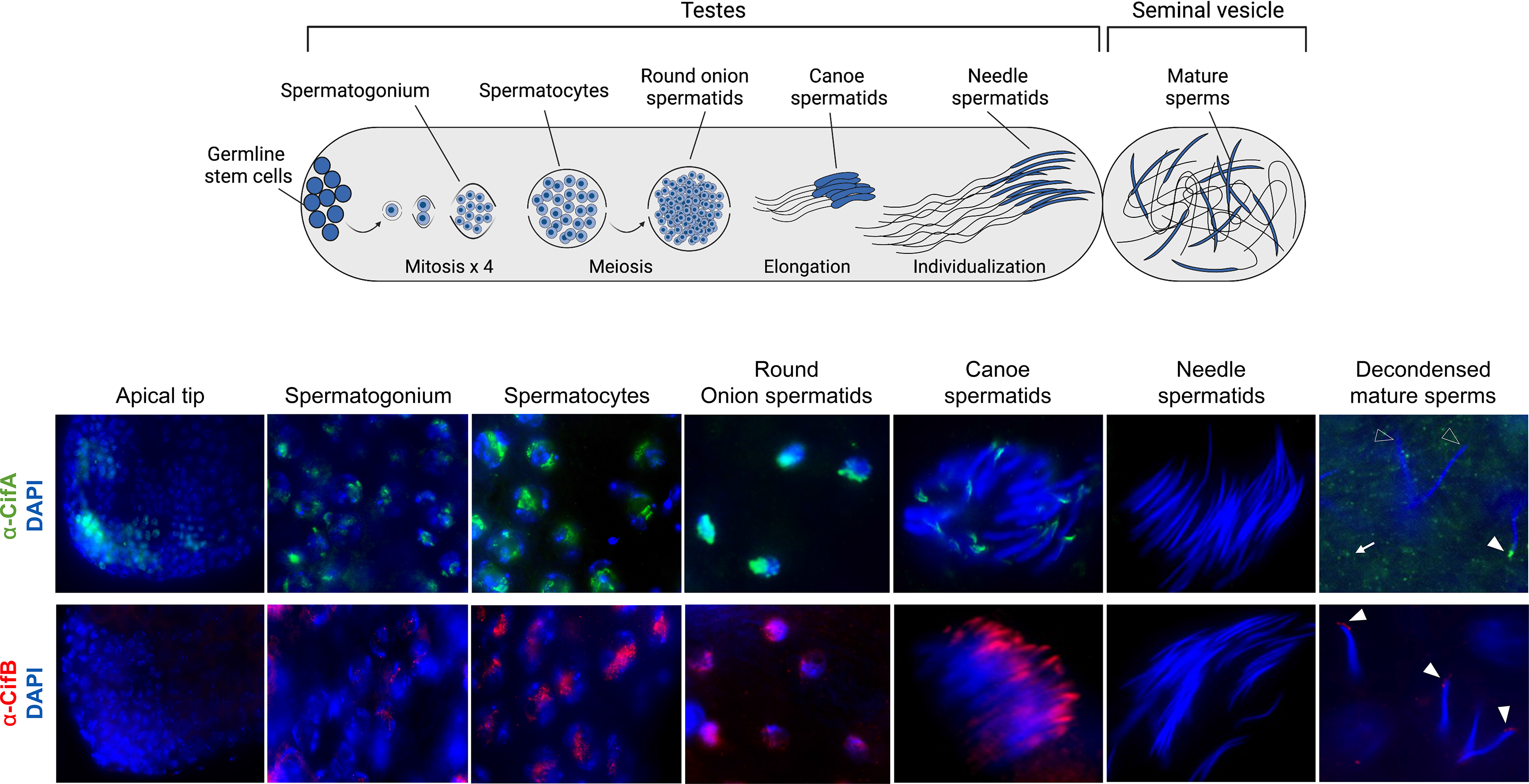
CifA and CifB invade sperm nuclei during spermatogenesis and spermiogenesis. Schematic representation of *Drosophila melanogaster* male reproductive system created by Biorender is shown on the top. Testes (n=20) from <8 hrs old males expressing dual transgenes *cifAB* were dissected and immunostained to visualize CifA (green) and CifB (red) during sperm morphogenesis. DAPI stain (blue) was used to label nuclei. CifA, but not CifB, localizes in the germline stem cells at the apical end of testes. Both CifA and CifB localize in the nuclei of mitotic spermatogonium, spermatocytes, and round onion stage spermatids. In the later stages of spermiogenesis, elongating spermatids harbor CifA and CifB at the acrosomal tip of the heads. CifB is present in all canoe-stage spermatid nuclei, whereas CifA is present on average in 39% of spermatids per bundle. Cifs are not accessible by the antibodies in the tightly compacted spermatids at the needle-stage. After decondensing mature sperms isolated from seminal vesicles (see methods), CifA and CifB are detectable in the acrosome regions at varying percentages. CifA is common among sperm tails in a speckled pattern (white arrow) and either present on average in 45% or 0% of the mature sperm heads depending upon the sampled seminal vesicles. CifA’s presence in the acrosome region is shown by solid white arrowheads and absence with empty white arrowheads. CifB is present in acrosomal tips of all of the sperms (solid white arrowheads) and does not occur with sperm tails. CifA and CifB localization patterns are similar in wild type (*w*Mel+) line and signals are absent in *Wolbachia*-uninfected (*w*Mel-) negative control line (Fig S2). Some of the CifA and CifB proteins are also stripped by the individualization complex into the cytoplasmic waste bag (Fig S5). The experiment was repeated in two biological replicates.

During spermatogenesis, *Wolbachia* are stripped into the cytoplasmic waste bags, which eliminate excess cytoplasmic material during the process of spermatid elongation [23]. Here, we show that some CifA and CifB proteins are also stripped by the individualization complex into the cytoplasmic waste bag (Fig S5). Since *Wolbachia* are not present in the mature sperms [36, 37], these data suggest that the Cif proteins exit *Wolbachia* cells during spermatogenesis to possibly interact with and modify sperm DNA (see below). Taken together, these findings demonstrate CifA and CifB proteins access *Drosophila* sperm nuclei throughout development.

### Cifs cause abnormal histone retention and protamine deficiency

Since the Cifs localize to developing sperm nuclei during spermatogenesis, we hypothesized that they may incipiently interact with nuclear DNA to impact sperm genome integrity – a central prediction of the Host Modification model of CI [11, 29]. At the histone-to-protamine transition stage during spermatogenesis [38], histones normally undergo various post-translational modifications (PTMs) for removal and replacement by smaller protamines for tight chromatin reorganization [38–41]. Lack of PTMs can lead to histone-bound chromatin with improper protamine deposition that causes paternal chromatin defects, male infertility, and embryonic lethality [27,42,43]. Thus, the incipient defects initiated in the testes can lead to post-fertilization catastrophes.

Utilizing a core histone antibody, we investigated histone abundance within spermatid bundles at the late canoe stage in CI- and non-CI causing males. We detected significantly increased histone retention in both *w*Mel+ and *cifAB*-expressing testes from <8 hr old males compared to the negative controls (Fig 2A, Fig S6, Table S1). Single transgenic expression showed significantly less histone-retaining bundles at this stage similar to *w*Mel- negative controls (Fig S7A, Table S1). To detect if abnormal histone retention is linked with protamine deficiency in mature sperms, we next used the fluorochrome chromomycin A3 (CMA3) stain that fluoresces upon binding to protamine-deficient regions of DNA [44, 45]. Mature sperms isolated from wild-type seminal vesicles of young (high CI-inducing) *w*Mel+ males exhibit increased protamine deficiency relative to *w*Mel- males (Fig 2B, Table S1).

**Fig 2.**
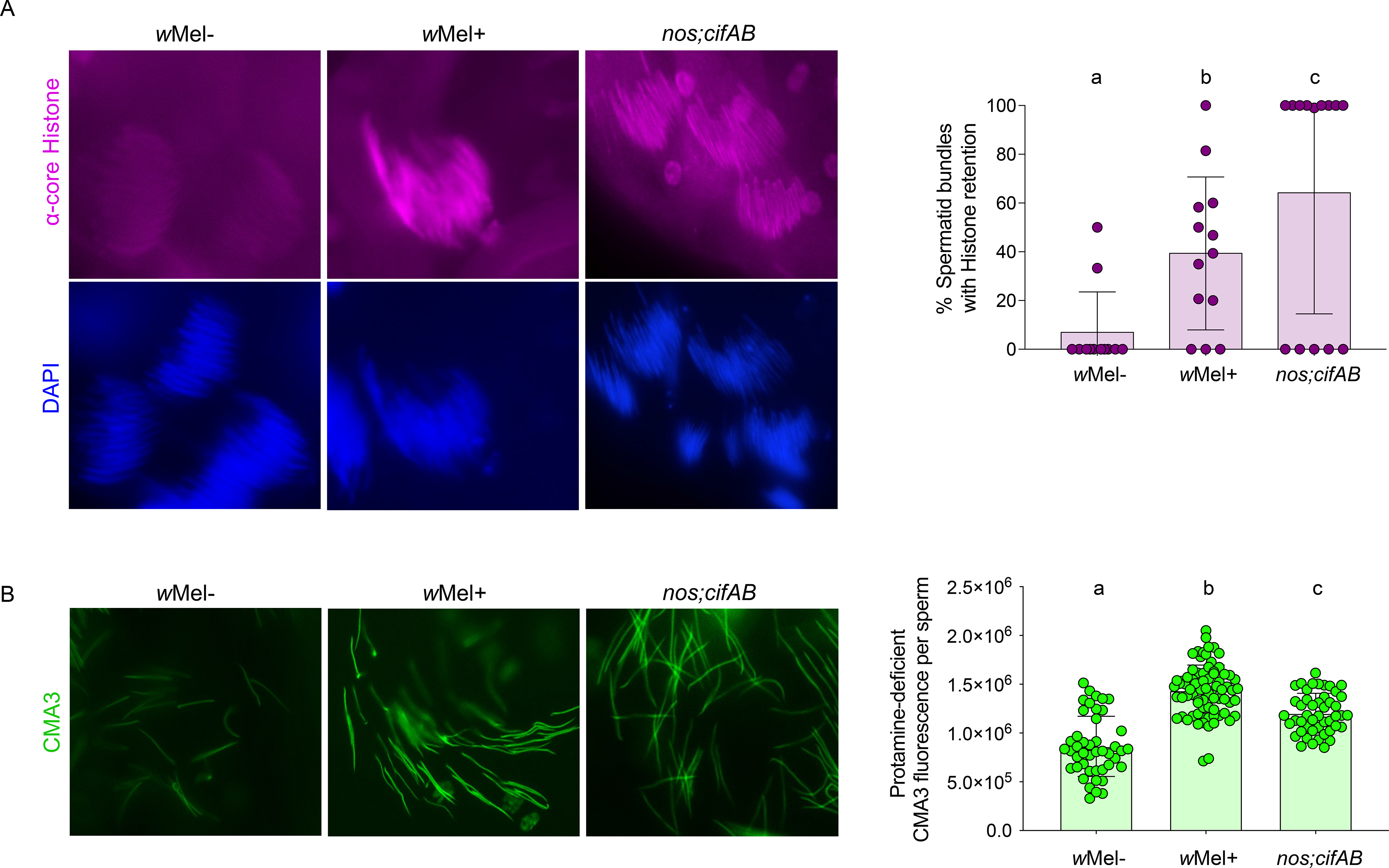
Cifs cause histone retention in late canoe spermatids and protamine deficiency in mature sperms. (A) Testes (n=15) from <8 hrs old males of *w*Mel+, *w*Mel- and transgenic *cifAB* lines were dissected and immunostained to visualize and quantify spermatid bundles with histone retention (purple) during late canoe stage of spermatogenesis. DAPI stain (blue) was used to label spermatid nuclei. Total spermatid bundles with DAPI signals and those with retained Histones were manually counted and graphed. Compared to the negative control *w*Mel-, *w*Mel+ *Wolbachia* and dually-expressed *cifAB* transgenic lines show abnormal histone retention in the late canoe stage. Vertical bars represent mean, and error bars represent standard deviation. Letters indicate statistically significant (p<0.05) differences as determined by pairwise comparisons based on Kolmogorov-Smirnov test. (B) Mature sperms isolated from seminal vesicles (n=15) of <8 hr old males reared at 21°C were stained with fluorescent CMA3 (green) for detection of protamine deficiency in each individual sperm nucleus. Individual sperm head intensity was quantified in ImageJ (see methods) and graphed. *w*Mel+ and transgenic *cifAB* lines show enhanced protamine deficiency levels compared to *w*Mel- control. Vertical bars represent mean and error bars represent standard deviation. Letters indicate statistically significant (p<0.05) differences as determined by multiple comparisons based on a Kruskal-Wallis test and Dunn’s multiple test correction. All of the P-values are reported in Table S1. The experiments were performed in two independent biological replicates and samples were blind-coded for the first run. Raw data underlying this figure can be found in S1 data file.

To investigate if lack of protamines associates with CI, we isolated sperms from <8 hr old males with a protamine A and B knockout mutant line. We show that protamine mutants, both in the presence (Prot+) and absence (Prot-) of *Wolbachia,* also exhibit a significant increase inΔ Δ fluorescence relative to *w*Mel- (Fig 3A, Table S1). Moreover, a key outcome of higher protamine deficiency in Prot+ males with *Wolbachia* is an increase in CI compared to wild type *w*Mel+ Δ CI, under the same experimental setup (Fig 3B, Table S2). These findings indicate an additive effect by *Wolbachia* and ΔProt knockouts on the protamine deficiency and CI penetrance.

**Fig 3.**
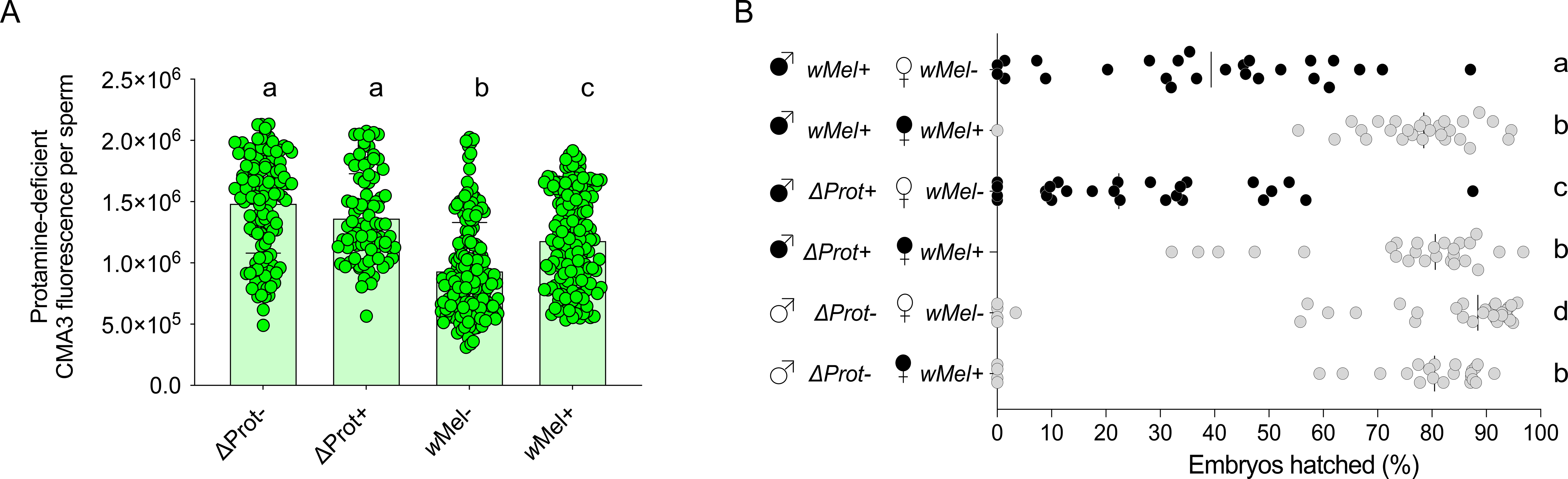
Protamine mutants enhance wild type CI and show significantly increased levels of protamine deficiency in mature sperms. (A) Sperms from the *Wolbachia*-infected (Prot+) and -uninfected (Prot-) protamine mutant (w[1118]; Δ Δ st35B[floxed], Sco/CyO) males exhibit significantly increased CMA3 fluorescence indicative of protamine deficiency compared to both wild type *w*Mel+ and *w*Mel-. Vertical bars represent mean, and error bars represent standard deviation. Letters indicate statistically significant (p<0.05) differences as determined by multiple comparisons based on a Kruskal-Wallis test and Dunn’s multiple test correction. All of the P- values are reported in Table S1. (B) CI hatch rate analyses of male siblings used in CMA3 assays (panel A) validate that Δ t+ males with increased sperm protamine deficiency causes stronger Pro (rescuable) CI levels than *w*Mel+. Δ rot- males do not cause CI. Letters to the right indicate statistically significant (p<0.05) differences as determined by pairwise Mann-Whitney test and multiple comparisons calculated using a Kruskal-Wallis test and Dunn’s multiple test correction. All of the P-values related to CI assay are reported in Table S2. Raw data underlying this figure can be found in S1 data file.

Notably, Δ rot- males do not recapitulate CI on their own; thus, the protamine deficiency is not P the sole cause of CI and must operate in conjunction with other CI modifications. Consistent with these results, 7-day old *w*Mel+ males that express almost no CI exhibit a similarly weak protamine deficiency level to *w*Mel- males, as expected (Fig S8A, Fig S8B and Table S2). Moreover, transgene analyses specify both single and dual expression of CifA and CifB cause protamine deficiencies at significantly higher levels than negative controls of *w*Mel- and a non-CI transgene (Fig S7B, Table S1). Since singly expressed Cifs do not cause CI in *D. melanogaster* [16] (Fig S7C, Table S2), additive effects on the protamine deficiency and/or histone retention due to abnormal PTMs may be required to fully establish CI.

### Both CI and rescue are dependent upon a CifA bipartite nuclear localization signal

Based on the cNLS mapping tool for nuclear localization signals [46], CifA amino acids harbor a predicted bipartite nuclear localization sequence (bNLS) (Table S3) in the most conserved region of the protein [13,47,48] that is under strong purifying selection [17]. As nuclear localization signals bind to the extended surface groove of nuclear transport protein importin-α, also known as karyopherin-α [49], we hypothesized that sperm nuclear localization of CifA and CI and rescue are dependent on the bNLS. To test this hypothesis, we mutagenized two bNLS sequences with alanine substitutions (aa189-190 for NLS1 (denoted *cifA_189_*), aa224- 225 for NLS2 (denoted *cifA_224_*)), and we additionally deleted the entire bNLS region (*cifA_ΔbNLS_*) (Fig 4A). The bNLS deletion also corresponds to the weakly predicted catalase-rel domain in CifA [47, 48].

**Fig 4.**
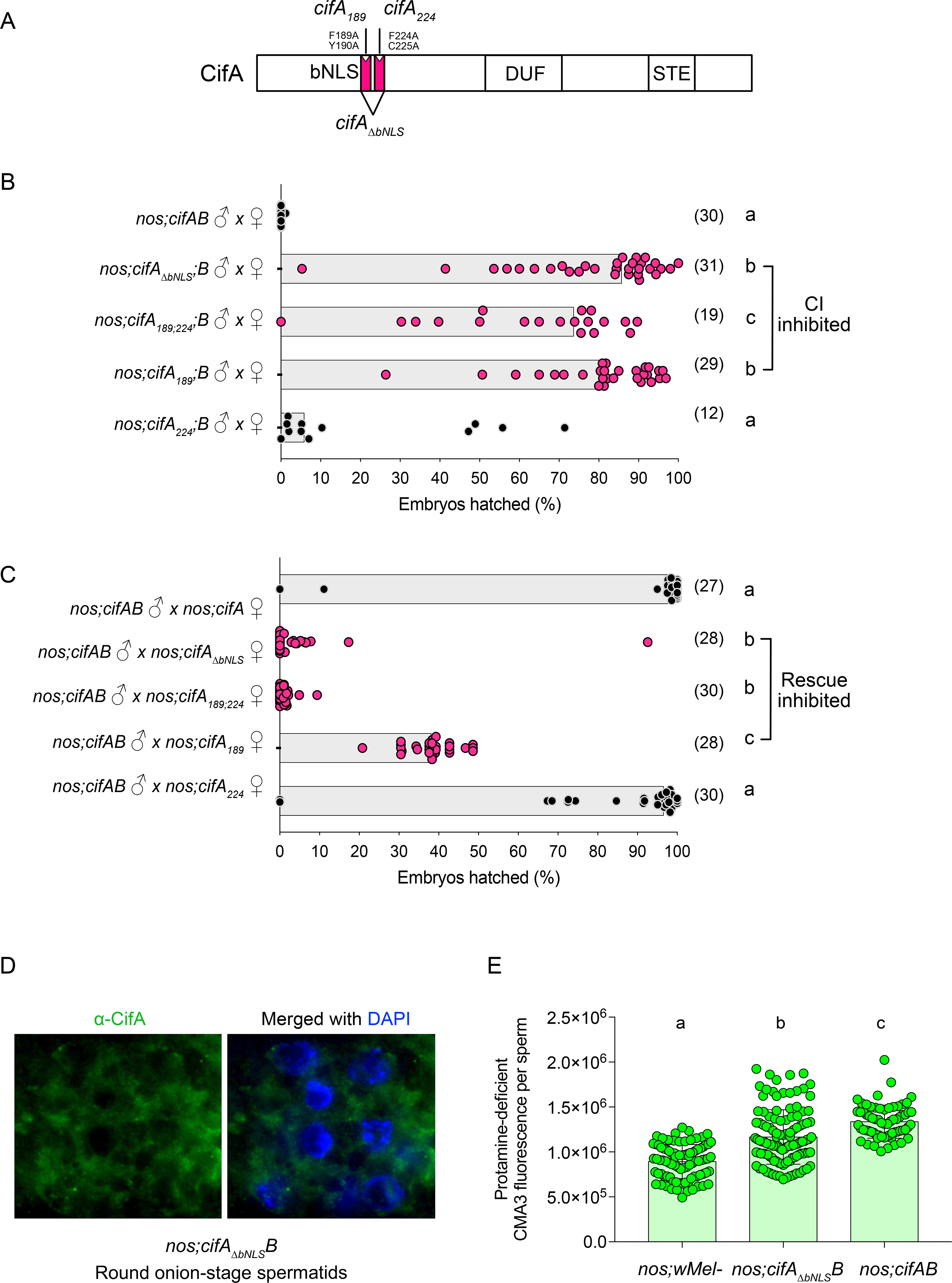
A nuclear localization signal in CifA is necessary for CI, rescue, and protamine levels. (A) Schematic representation of CifA annotation shows the annotated bipartite nuclear localization signal (bNLS) with engineered amino acid substitutions and deletions. (B, C) Hatch rate assays assessed both CI (B) and rescue (C) in flies expressing wild type, transgenic, and mutant *cifA*. Each dot represents the percent of embryos that hatched from a single male and female pair. Sample size is listed in parentheses. Horizontal bars represent the median. Letters to the right indicate significant differences determined by a Kruskal-Wallis test and Dunn’s multiple comparison tests. All the P-values are reported in Table S2. (D) Antibody labeling (green) and DAPI staining of onion stage spermatids in the testes of the bNLS mutant line (*cifA_⊗bNLS_*) reveals that the deletion ablates CifA’s localization to the nucleus, and CifA thus remains in the surrounding cytoplasm. The imaging experiment was conducted in parallel to *nos;cifAB* line shown in Fig 1. (E) Mature sperms isolated from seminal vesicles (n=15) of <8 hr old males of transgenic *cifA _bnls_B* line shows reduced fluorescence indicative of less Protamine deficiency compared to *cifAB*. To control for any background confounding effects of *nos- Gal4VP16* driver line, *w*Mel- fathers were prior crossed to *nos-* mothers to generate males with *nos;wMel-* genotype. CMA3 fluorescence levels of sperms isolated from *nos;wMel-* males were similar to *w*Mel- wild type lines used in previous assays in this study. Vertical bars represent mean and error bars represent standard deviation. Letters indicate statistically significant (p<0.05) differences as determined by multiple comparisons based on a Kruskal-Wallis test and Dunn’s multiple test correction. All of the P-values are reported in Table S1 and raw data underlying this figure can be found in S1 data file.

Each bNLS mutant, individually and together (*cifA_189;224_*), was dually expressed in testes with transgenic, intact *cifB* to assess CI and singly expressed in females to assess rescue. Transgenic *cifA_189_* expression significantly reduced CI and rescue as previously reported (Fig 4B,C and Table S2) [48]. Conversely, transgenic *cifA_224_* expression showed no significant difference from the controls in either CI or rescue, suggesting this region has little to no impact. However, when both mutants are expressed in *cifA_189;224_* or when the entire bNLS is deleted, CI and rescue are strongly inhibited (Fig 4B,C and Table S2). These results highlight the importance of the nuclear localization sequence in inducing CI as well as rescue. To determine if the lack of CI induction is due to non-nuclear localization of CifA protein, we used the deletion mutant *cifA_⊗bNLS_* with wild type *cifB* to demonstrate that in contrast to its normal, nuclear localization (Fig 1), it mislocalizes to the cytoplasm of onion stage spermatids rather than the nuclei (Fig 4D). Additionally, to test if deletion of the bNLS impacts sperm genomic integrity, we performed CMA3 based protamine-deficiency assay (as described above) and found reduced protamine deficiency levels in matures sperms due to non CI-causing *cifA_ΔbNLS_;B* line compared to CI-causing *cifAB* (Fig 4E, Table S1), providing further evidence that protamine deficiency is linked to CI. Overall, these data provide previously unknown findings that a functional nuclear localization sequence and CifA nuclear localization impacts CI, rescue, and sperm protamine levels.

### Cifs cause a paternally-transferred protamine deficiency

To investigate if Cif proteins and/or the genome integrity modifications transfer paternally to the female reproductive tract, single male:female pairwise matings were set up for CI and rescue crosses. After 4 hrs of mating, we isolated the whole uterus (Fig 5A) including the sperm storage organs (spermathecae (SP) and seminal receptacle (SR)). Following sperm decondensation, antibody staining, and microscopy, we observed two key results. First, both CifA and CifB proteins transfer with sperm tails and heads, respectively, to the sperm storage organs of *Wolbachia*-free females (Fig 5B). Second, the CI-associated sperm protamine deficiency induced by *w*Mel+ and *cifAB*-expressing males transfers and persists in the sperms isolated from SP and SR of *Wolbachia*-free, mated females (Fig 5C,D and Table S1). These findings connect a paternally-transferred sperm modification with the activity of *Wolbachia* and the Cifs themselves. Results strongly support the Host Modification model of CI since a Cif- induced sperm modification established in the testes transfers to the female reproductive tract.

**Fig 5.**
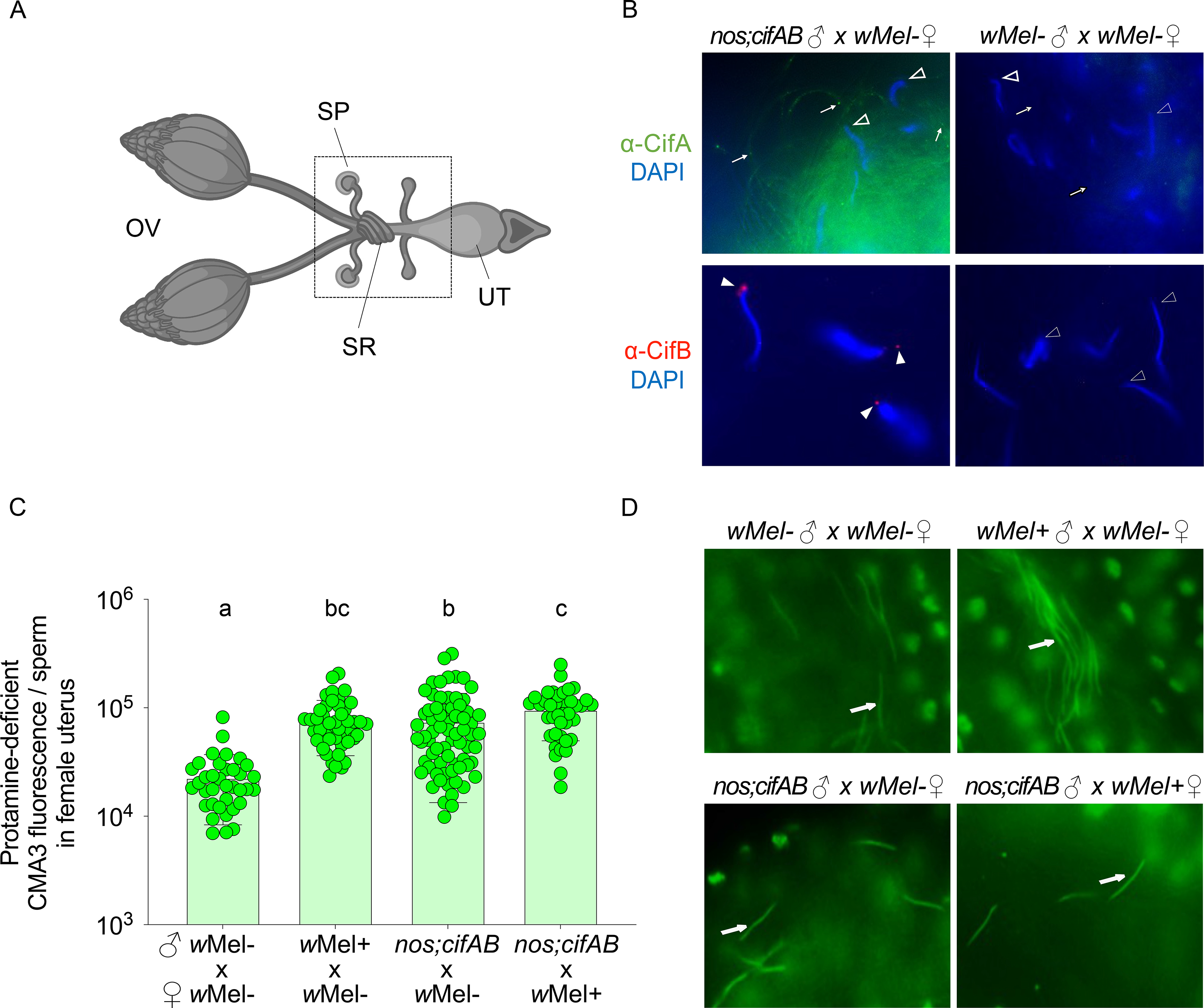
CifA, CifB, and the protamine deficiency are transferred with the mature sperm to the female reproductive tract. (A) Schematic representation of *Drosophila melanogaster* female reproductive system. Mature oocytes leave the ovary (OV) and reach the uterus (UT), where they can be fertilized prior to being laid. Sperms from males are stored in specialized organs - spermathecae (SP) and seminal receptacle (SR) shown in the box, which open into the UT for fertilization to occur. Schematic is created with BioRender (B) Transgenic *cifAB*- expressing and *w*Mel- males were crossed to *w*Mel- females. 4 hrs post-fertilization, sperms isolated from females were decondensed and immunostained for localizing CifA (green) and CifB (red). DAPI stain (blue) was used to label nuclei. CifA is absent in sperm heads (empty arrowheads) and puctae are seen along the sperm tails (arrows). CifB is present in apical acrosomal tip of all of the sperm heads (solid arrowheads), with more distant signal in the more decondensed sperm nuclei. No Cifs are present in the sperms transferred from *w*Mel- negative control males. (C) Individual sperm intensity quantification shows that protamine deficiency of sperms from *w*Mel+ and transgenic *cifAB* males persists after transfer in the females compared to *w*Mel- males. Sperm protamine deficiency from transgenic *cifAB* males also persists in the reproductive tract of *w*Mel+ females. Vertical bars represent mean, and error bars represent standard deviation. Letters indicate statistically significant (p<0.05) differences as determined by multiple comparisons based on a Kruskal-Wallis test and Dunn’s multiple test correction. All of the P-values are reported in Table S1 and the raw data underlying this panel can be found in S1 data file. (D) Representative images of CMA3-stained mature sperms (arrows) transferred from *w*Mel-, *w*Mel+ and transgenic *cifAB* males in *w*Mel- and *w*Mel+ female reproductive systems are shown.

### CifA is present in early oogenesis and absent from mature eggs and rescue embryos

Expression of *cifA* alone in the ovaries rescues CI [16, 17], yet how CifA protein mediates rescue is unknown and central to further differentiating the mechanistic models of CI. For instance, CifA in females may modify reproductive cell biology to nullify CI-inducing sperm modifications in the embryo (Host-Modification), or alternatively, CifA may directly bind CifB in the embryos to prevent its proposed CI toxicity (Toxin-Antitoxin). Using CifA antibodies, we show in *w*Mel+ and *cifA* transgene-expressing ovaries that CifA protein is cytonuclear and specifically localizes to cyst DNA in region 1 of the germarium (Fig 6A), indicative of nuclear access in ovaries similar to that in testes (Fig 1). Cystoblast in the germarium undergoes rounds of mitotic divisions to produce oocyte and nurse cells [50, 51]. Along egg chamber stages 2-8 of *w*Mel+ females, CifA colocalizes with *Wolbachia* in the nurse cells and oocyte cytoplasm (Fig 6A). While *Wolbachia* are abundant in the stage 10 egg chamber, CifA is notably absent. In transgenic females, CifA is also primarily detected in the germarium and cytoplasm of the early egg chambers and absent in late egg chambers (Fig 6A, Fig S9A). Presence of high levels of CifA in *Wolbachia*-infected eggs is proposed to rescue CI; importantly, we did not detect CifA in ∼30-60 min old rescue embryos during early mitotic divisions (Fig 6B). Moreover, CifA was not detected in 1-2 hr old embryos (Fig S9B), whereas the positive control histone signals colocalize with embryonic DNA at both embryonic developmental stages.

**Fig 6.**
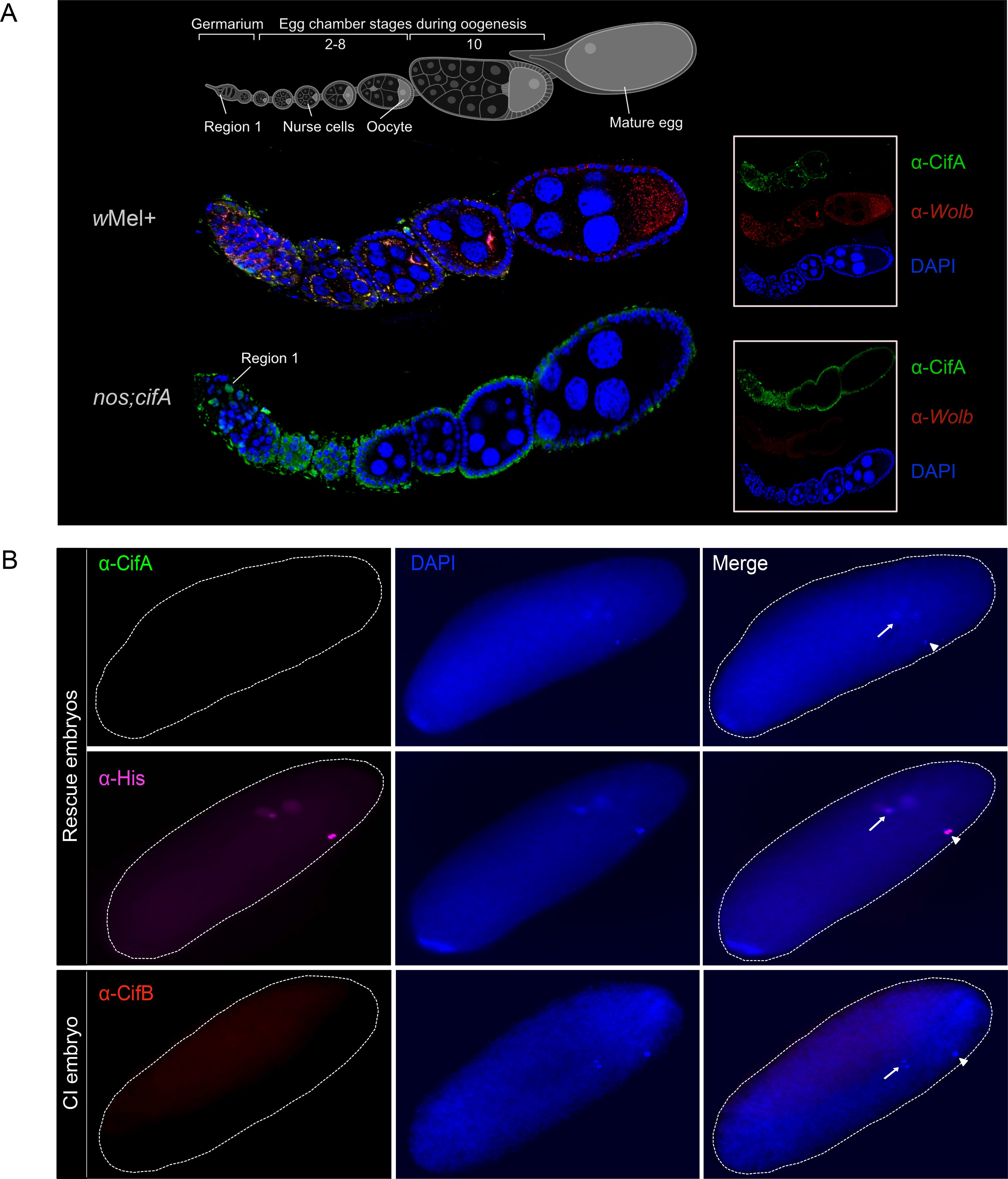
CifA is present in early oogenesis and absent in late-stage egg chambers. Both CifA and CifB are absent in CI and rescue embryos. (A) Schematic representation of *Drosophila melanogaster* ovariole at the top illustrates the stages of oogenesis from left to right. Image was created with BioRender. Immunostaining assay indicates localization of CifA (green) to the cyst DNA (blue labeled with DAPI) in region 1 of the germarium of *Wolbachia*-uninfected transgenic *cifA* line. In *w*Mel+ line, CifA colocalizes with *Wolbachia* (red) in the germarium, nurse cells and oocyte cytoplasm along 2-8 stages of egg chambers. CifA is absent in stage 10 egg chamber, whereas *Wolbachia* signals persist. In the transgenic *cifA* line, we note the observed autofluorescence in green channel outlining the tissue morphology does not signify CifA signals. Ovariole images were manually adjusted in Affinity designer software to align egg chamber stages in the same plane. (B) Immunofluorescence of CifA (green) and CifB (red) in ∼30-60 min old embryos obtained from rescue (*cifAB* male x *w*Mel+ female) and CI crosses (*nos;cifAB* male x *w*Mel- female). Histone antibody labeling core-histones (magenta) was used as a positive control. Histone signals were detected colocalizing with host DNA, labelled with DAPI (blue), whereas no CifA and CifB signals were detected. Dotted white embryonic periphery is drawn around the embryo shape. White arrows indicate dividing nuclei post fertilization and arrowheads indicate polar bodies.

CifB from *w*Pip was recently shown to be paternally-transferred to the CI embryos [30]. We evaluated if *w*Mel CifB colocalizes with mitotic, embryonic DNA after fertilization. CifB is absent from embryonic nuclei and the cytoplasm of CI embryos (Fig 6B), suggesting that CifB is not inherited with the paternal DNA to the embryo. The absence of CifA and CifB in late wild type and transgenic eggs and embryos indicates that host gametic changes prime the embryo for CI and rescue before fertilization, as predicted by the Host Modification model of CI. This inference is also consistent with previous studies where ovarian, rather than embryonic, transgenic *cifA* expression rescues CI [13,15–17,52].

## Discussion

At the genetic level, dual expression of *cifA and cifB* or single expression of *cifB* can recapitulate CI; and *cifA* rescues CI [13,15–19,52]. However, the cellular and mechanistic bases of CI and rescue remain unresolved and the subject of several questions: When and where do the Cif proteins localize in testes to potentiate CI? Are the Cifs transferred to the embryo already modified for CI-defining defects? Do CifA and CifB bind in the embryo to rescue lethality? Here, we establish that both CifA and CifB proteins invade nuclei of developing spermatids and modify paternal genome integrity by altering the normal Histone-Protamine transition process. Specifically, dual CifA and CifB expression induces abnormal histone retention and protamine deficiency in CI-causing male gametes to induce CI. Moreover, knocking out protamines enhances wild type CI, and a nuclear localization sequence in CifA is essential for CI, rescue, and protamine deficiency. Finally, binding of CifA and CifB in the embryo is not evident.

Sperm genome compaction is normally achieved during the post-meiotic canoe phase of spermatogenesis, when histones are replaced by protamines [53]. This compaction process is highly conserved from flies to humans [54–56] and plays a crucial role in successful fertilization and embryonic development [57, 58]. Histone marks are carriers of transgenerational epigenetic information [59], and changes in the sperm epigenome can lead to detrimental consequences including early embryonic lethality and birth defects [59, 60]. Histones undergo various post- translational modifications (PTMs) such as ubiquitination, methylation, phosphorylation and acetylation before degradation and removal from the sperm chromatin [38]. Therefore, abnormally-retained histones could result from aberrant PTMs in Cif-expressing flies. In a protein interactome screen, both ubiquitin and histone H2B were determined as binding host candidates to CifB [52]. Thus, it is possible that inhibition of the histone ubiquitination process mediates abnormal histone retention. Additionally, histone acetylation during the canoe stage of spermatogenesis is required for histone eviction in *Drosophila* [61]. Indeed, reduced acetylation levels lead to abnormal histone retention and protamine deficiency, which causes embryonic inviability in flies [61] and in mammals [40,62,63]. Future research investigating what specific PTMs are altered at the Histone-Protamine transition stage will help elucidate the molecular pathway(s) leading to CI.

The paternally-derived Cif proteins travel with the sperms to the female reproductive tract, where CifB is present in the acrosomal region and CifA occurs along the tail. Notably, both Cifs are not evident in the embryos. While the presence of paternally-transferred Cifs is ambiguous to the mechanistic models of CI [11, 29] and recently confirmred in another transgenic study [30], it is the presence of CifA-CifB binding in the rescue embryo that would support the Toxin-Antitoxin model. However, there is no evidence of this binding phenomenon in the embryos to date. Thus, we conclude that Cifs act before fertilization to prime the sperm chromatin and incipiently launch CI. Paternal effect proteins can modify sperm in various systems to bestow embryonic defects, even though the proteins themselves do not transfer to the embryos [60,64,65].

In *Drosophila*, the sperm enters the egg with an intact membrane [66]. Therefore, absence of Cifs in the embryos raises the question at what point the presence of CifA along the tail and CifB in the acrosome is lost before the sperm enters the egg. One possible explanation is that the Cif proteins are released from the sperm upon exocytosis of the acrosome. The acrosome, best known as a secretory vesicle, undergoes exocytosis and releases its contents to facilitate sperm-egg binding in mammals [67, 68]. Though not well characterized in insects, studies in the house fly *Musca domestica* suggest loss of the sperm plasma membrane before entry into the egg, followed by exocytosis of acrosomal contents during passage of the sperm through the egg micropyle [69].

CifA is absent in *Wolbachia*-infected and transgenic embryos, which indicates that rescue is established during oogenesis under the Host Modification model, and CifA thus does not bind and nullify CifB from *w*Mel *Wolbachia* in the embryos. Indeed, a CifA mutant in the newly annotated nuclear localization sequence ablates rescue, suggesting access to ovarian nuclei is important for rescue. Interestingly, structures of CifA and CifB support binding of the two proteins [70], yet in light of these results here, the CifA-CifB binding may be central to CI induction instead of rescue. Moreover, mutating amino acid sites across the length of the CifA protein, including binding and non-binding residues, generally ablates CI and rescue [48, 70]. Thus, it is possible that CifA mutants that ablate rescue do so by altering the protein structure, function, and/or location of ovarian targets to modify, rather than the embryonic binding of CifA to CifB *in vivo*. Future work will be important to resolve how CifA primes oogenesis to alter specific cell biological and biochemical events that underpin rescue.

Once fertilization occurs, protamines from the paternal chromatin must be removed and replaced by maternal histones to decondense and activate the chromatin of the developing embryo [71]. Interestingly, post-fertilization delays in maternal H3.3 histone deposition occur in CI embryos [72]. The delay may in part be due to pre-loaded paternal histones or altered paternal epigenome information leading to mistiming of maternal histone deposition, hence causing CI. Thus, we propose that a genome integrity network involving histones, protamines, and possibly other factors in the gametes may be a common and defining feature underpinning the onset of CI and rescue.

Altogether, discovery of nuclear-targeting Cif proteins in male and female gametes establishes new insights on the early cell biological and biochemical steps that underpin the CI drive system with relevance to arthropod speciation and pest control [11]. In addition to disentangling the reproductive events of the Cif proteins that control gametogenesis and embryogenesis, the evidence specifies that the Cif proteins modify sperm genomic integrity and transfer paternally, but they themselves do not enter and bind each other in the embryo to enable rescue. These findings are consistent with the Host Modification model of CI by *w*Mel *Wolbachia*. More generally, as there are no previous reports of prophage proteins invading animal gametic nuclei to impair the histone-protamine transition during spermatogenesis, the findings have implications for expanding an appreciation of prophage-bacteria-eukaryote interactions into the realm of animal reproduction.

## Material and Methods

### Cif proteins antibody development

Conserved amino acid regions of CifA and CifB proteins from *w*Mel *Wolbachia* were previously identified [47]. Using these regions, monospecific polyclonal antibodies were commercially generated by Pacific Immunology through injection of three synthesized and conserved short (20 aa) peptides for each protein into rabbits. Sequences of peptides were Cys- EYFYNQLEEKDKEKKLTE for CifA, and Cys-DENPPENLLSDQTRENFRR for CifB. The resulting *α*-CifA and *α*-CifB antibodies were evaluated using an enzyme-linked immunosorbent assay, and titers were determined to be higher than 1:500,000 for each antibody. Using standard protocols of the Invitrogen WesternDot kit, antibody specificity to *w*Mel+ samples was verified using western blots (1:1000-fold antibody dilution) on protein isolated from homogenates of 50 testes pairs (0-8 hour old males) and 10 ovary pairs (6 day old females) from *w*Mel+ (positive), *w*Mel- (negative control), and *cifAB* transgenic (positive) flies. The correct size band was only detected from *w*Mel+ and *cifAB* reproductive tissues (Fig S1). Because the antibodies were generated in the same animal, all subsequent labeling was done with individual antibodies.

### NLS identification

CifA amino acid sequences from known *Wolbachia* and close relatives were input into the cNLS Mapper software [73] to identify putative NLS sequences within each protein (Table S3). cNLS Mapper identifies sequences specific to the importin α/β pathway. A cut-off score of 4 was applied to all sequences. Higher scores indicate stronger NLS activities. Scores >8 indicate exclusive localization to the nucleus, 7-8 indicate partial localization to the nucleus, 3-5 indicate localization to both the nucleus and the cytoplasm, and score 1-2 indicate localization exclusively to the cytoplasm. Predicted NLS sequences are divided into monopartite and bipartite classes. Monopartite NLSs contain a single region of basic residues, and bipartite NLSs contain two regions of basic residues separated by a linker region.

### Development of transgenic lines

A *cifA* variant was synthesized de novo at GenScript and cloned into a pUC57 plasmid as described previously [48]. Site-directed mutagenesis was performed by GenScript to produce the mutants outlined in Figure 5. The *cifA_189_* variant was first described in Shropshire et al. [48] as *cifA*_2_. UAS transgenic *cifA* mutant flies were then generated using previously established protocols [13]. Briefly, GenScript sub-cloned each gene into the pTIGER plasmid, a pUASp- based vector designed for germline-specific expression. Transgenes were then integrated into y^1^ M{vas-int.Dm}ZH-2A w*; P{CaryP}attP40 attachment sites into the *D. melanogaster* genome using PhiC31 integrase via embryonic injections by BestGene. At least 200 embryos were injected per transgenic construct, and successful transformants were identified based on red eye color gene included on the pTIGER plasmid containing the transgene. All sequences are reported in Table S4.

### Fly rearing and strains

*D. melanogaster* stocks y^1^w* (BDSC 1495), *nos*-GAL4:VP16 (BDSC 4937), UAS transgenic lines homozygous for *cifA*, *cifB*, *cifAB, WD0508* [13] and Protamine mutant (w[1118]; ΔMst35B[floxed], Sco/CyO) [74] were maintained on a 12-hour light/dark cycle at Δ25^0^C and 70% relative humidity on 50mL of standard media. Uninfected protamine mutant line was generated by three generations of tetracycline treatment (20 μ/ml in 50 ml of fly media) as described in previous studies [13], followed by two rounds of rearing on standard food media before using in the experiments. Infection status for all lines was regularly confirmed by PCR using Wolb_F and Wolb_R3 primers [75].

### Hatch rates

Parental flies were either wild type uninfected (*w*Mel-) or infected (*w*Mel+) with *Wolbachia* or transgene-expressing with no *Wolbachia* infection. Uninfected transgenic flies were generated previously [13, 17]. Paternal grandmother age was controlled to 9-11 days for expression of naturally high penetrance of *w*Mel CI [76]. Parental transgenic males were generated through crossing *nos*-Gal4:VP16 virgin females (aged 9-11 days) to UAS-*cif* transgenic, uninfected males [76]. Mothers were aged 6-9 days before crossing, while father males first emerged between 0-8 hours were used in hatch rates and tissue collections to control for the younger brother effect associated with lower CI penetrance [13, 77].

Hatch rates were set up as described previously [13, 17]. Briefly, a male and female pair was placed in an 8oz, round bottom, polypropylene *Drosophila* stock bottle with a grape juice-agar plate containing a small amount of yeast placed at the base and secured with tape. These bottles were then placed in a 25°C incubator overnight to allow for courting and mating. The following day, these plates were discarded and replaced with new grape juice-agar plates with fresh yeast. After an additional 24 hours, the plates were removed, and the embryos were counted. The embryo plates were then incubated for 36 hours at 25°C before the total number of unhatched embryos were counted. Any crosses with fewer than 25 embryos laid were discarded from the analyses. Statistical significance (p<0.05) was determined by a Kruskal-Wallis test and Dunn’s multiple test correction in GraphPad Prism 7. All p-values are listed in Table S2.

### Immunofluorescence: testes and seminal vesicles

Siblings from the hatch rate (males 0-8 hours) were collected for testes dissection in ice-cold 1x PBS solution. Tissues were fixed in 4% formaldehyde diluted in 1x PBS for 30 min at room temperature and washed in 1x PBS-T (1x PBS + 0.3% TritonX-100) three times for 5 min each. Tissues were then blocked in 1% BSA in PBS-T for 1 hour at room temperature. They were then incubated with 1° antibody (α-CifA 1:500 OR α-CifB 1:500) overnight at 4°C rotating. After washing in 1x PBS-T three times for 5 min each at room temperature, they were incubated with 1:1000 dilution Goat anti-rabbit Alexa Fluor 594 secondary antibody (Fisher Scientific, Cat#A11037) for 4 hours at room temperature in the dark. Tissues were then washed three times for 5 minutes each in 1X PBS-T and mounted on slides. To stain the nuclear DNA, 0.2mg/mL of DAPI was added to the mounting media before the coverslip was gently placed over the tissue and excess liquid wiped away. Slides were allowed to dry overnight in the dark before viewing on the Zeiss LSM 880 confocal microscope. All images were acquired with the same parameters for each line and processed in ImageJ as described in [78].

### Decondensation of Mature Sperm Nuclei

Squashed seminal vesicles collected from male flies (aged 0-8 hours) were treated with 10 mM DTT, 0.2% Triton X-100, and 400 U heparin in 1X PBS for 30 min [34]. The slides were then washed quickly in 1X PBS before immunofluorescence staining (see above).

### Immunofluorescence and quantification: Histones

Testes from male flies (aged 0-8 hours) were fixed and stained as described above for testes. The tissues were stained with a core histone antibody (Millipore Sigma, Cat#MABE71) (1:1000) and imaged on a Keyence BZ-800 microscope. Total late canoe stage sperm bundles were quantified in each testis, and those that retained histones were determined. Ratios of late canoe stage bundles containing histones relative to total bundles from each individual testis were graphed in GraphPad Prism 7. Statistical significance (p<0.05) were determined by pairwise comparisons based on Kolmogorov-Smirnov test and multiple comparisons based on a Kruskal-Wallis test and Dunn’s multiple test correction in GraphPad Prism 7.

### Sperm isolation and chromomycin A3 staining/quantification

Seminal vesicles were collected from male flies (aged 0-8 hours for 1-day old flies and 7- days for older flies) and placed on a microscope slide in ice-cold 1x PBS. Sperm was extracted on the slide using forceps and fixed in 3:1 vol/vol methanol:acetic acid at 4°C for 20 min. Excess solution was then removed, and the slide was air dried. Each slide was treated in the dark for 20 min with 0.25mg/mL of CMA3 in McIlvain’s buffer, pH 7.0, with 10mM MgCl2. Sperm was then washed in 1x PBS, mounted, and imaged using a Keyence BZ-X700 Fluorescence microscope. All images were acquired with the same parameters for each line and did not undergo significant alteration. Fluorescence quantification was performed by scoring fluorescent pixels in arbitrary units (A.U.) within individual sperm heads using ImageJ as per the details described in [78], and calculated fluorescence intensity per sperm head was graphed. Statistical significance (p<0.05) was determined by a Kruskal-Wallis test and Dunn’s multiple test correction in GraphPad Prism 7. All of the experiments involving CMA3 staining were performed at 21^0^C instead of 25^0^C. CI hatch rate assays were run in parallel to ensure that CI and rescue phenotypes are not impacted due to changed temperature conditions.

### Immunofluorescence: ovaries

Ovaries from females (6 days old) were dissected in 1x PBS on ice and processed as described previously [77, 79]. Tissues were blocked in 1% BSA in PBS-T for 1 hour at room temperature and were first incubated with α-CifA (1:500) primary antibody at 4^0^C overnight. After washing in 1x PBS-T three times for 5 min each at room temperature, they were incubated with 1:1000 dilution Alexa Fluor 488 secondary antibody (Thermo Fisher Scientific, Cat#A11034) for 4 hours at room temperature in the dark. Samples were then rinsed properly and blocked again before incubating with α-ftsZ (1:150) primary antibody (a kind gift from Dr. Irene Newton) to stain *Wolbachia* at 4^0^C overnight. After washing in 1x PBS-T three times, samples were incubated with second secondary antibody (Alexa Fluor 594) for 4 hours in the dark. Since both CifA and ftsZ antibodies were generated in the same animal, we used secondary antibodies conjugated to two distant fluorophores to distinguish specific signals. Tissues were then washed three times for 5 minutes each in 1X PBS, stained with DAPI to label nuclear DNA and mounted on slides. Slides were allowed to dry overnight in the dark before viewing on the Zeiss LSM 880 confocal microscope.

### Immunofluorescence: embryos

After 24 hours of mating, plates were switched, and embryos were collected every 30 minutes. Embryos were collected in a 100μm mesh basket in embryo wash solution. To remove the chorion, the basket was placed in 50% bleach for 3 min and then rinsed with 1x PBS. The embryos were then transferred to 50:50 4% paraformaldehyde (PFA) and heptane in a microcentrifuge tube and rotated for 20 min at room temperature. Tubes were then removed from the rotator, and the heptane and PFA were allowed to separate before the bottom PFA phase was carefully removed. Methanol was added to the remaining heptane, and the tube was shaken vigorously for 20 seconds before the embryos settled to the bottom and solution was removed. A new volume of methanol was added to the embryos, and they were allowed to settle to the bottom of the tube. Methanol was removed, and all blocking, staining, and imaging steps were carried out for testes and ovary tissues above.

## Acknowledgments

The authors thank Jennifer Battle for her assistance in fly collections and staining for sperm integrity assays, Alex Mansueto for his assistance in hatch rates, and Sarah Bordenstein, Dylan Shropshire, and Luis Mendez for providing helpful feedback on the manuscript. We thank Dr. Janna McLean for sending the Protamine mutant fly line for conducting the experiments. We thank Dr. Irene Newton for sharing *Wolbachia* antibody and providing useful feedback on the preprint version of this manuscript. We also thank the Cell Imaging Shared Resource (CISR) at Vanderbilt for imaging assistance.

## Author contributions

Conceptualization, B.A.L. and S.R.B.; Methodology, B.A.L., R.K. and S.R.B.; Investigation, B.A.L., R.K., and I.T.R.; Writing – Original Draft, B.A.L., R.K. and S.R.B.; Writing – Review & Editing, B.A.L., R.K., I.T.R. and S.R.B.; Funding Acquisition, B.A.L. and S.R.B.; Resources, S.R.B.; Supervision, S.R.B.

## Data and materials availability

All data are available in the main text or the supplementary materials. Unique biological materials will be available upon request.

## Supplementary figure legends

**Fig S1. Western blots using Cif antibodies reveal proteins at the proper size.** Western blots were run on protein extracted from ovaries (n=10) of wild type infected *w*Mel+, uninfected *w*Mel-, *cifA* transgenic (*cifA*), and dual *cifA;B* expressing transgenic lines. The expected size for CifA is ∼54 kD. Western blots were run using anti-CifB antibody on testes (n=15) of wild-type infected (+), uninfected (-), and *cifA;B* transgenic (A;B) lines. Expected CifB size is ∼133kD. Cifs are absent in *w*Mel- control and present at accurate size in *w*Mel+ and *cif expressing* lines.

**Fig S2. Cifs invade spermatid nuclei in wild type *w*Mel+ testes.** Testes (n=20) from <8 hrs old males of wild type *w*Mel+ and *w*Mel- lines were dissected and immunostained to visualize CifA (green) and CifB (red) during sperm morphogenesis. DAPI stain (blue) was used to label nuclei. CifA and CifB localization patterns in wild type lines are similar to that of transgenic *cifAB* (Fig 1) and signals are absent in *w*Mel- uninfected control line. The experiment was conducted in parallel to the one shown in Fig 1.

**Fig S3. CifA and CifB vary in abundance levels in spermatids and mature sperms.** (A) ImageJ-based signal intensity quantification indicates CifA (green) is more abundantly expressed than CifB (red) in the spermatogonium stage of the spermatogenesis. Mean of individual data points with standard deviation is plotted on the graph. Letters indicate statistically significant (p<0.05) differences as determined by pairwise comparisons based on a Mann-Whitney test. P-values are reported in Table S1. (B) In the decondensed mature sperms isolated from seminal vesicles, CifB is present in the acrosomal tip of canoe-shaped spermatids and mature sperm heads, whereas CifA is present in only 40% and 20% of them, respectively. Quantification was performed on the images obtained in Fig 1 data. Each dot represents percentage of Cifs present in spermatids or mature sperms per testes examined. Raw data underlying this figure can be found in S1 data file.

**Fig S4. CifA and CifB are not detectable in the condensed mature sperms in seminal vesicles due to technical limitations.** Seminal vesicles (n=20) from <8 hrs old males of transgenic *cifAB*, wild type *w*Mel+ and *w*Mel- lines were dissected and immunostained to visualize CifA (green) and CifB (red) in the mature condensed sperms (indicated by white arrows). DAPI stain (blue) was used to label nuclei. Absence of both CifA and CifB indicates that proteins are not accessible to the antibodies when the sperm chromatin is condensed and tightly packed.

**Fig S5. CifA and CifB are also removed in the cytoplasmic waste bag.** Testes (n=20) from <8 hrs old males of transgenic *cifAB*, and wild type *w*Mel- lines were dissected and immunostained to visualize CifA (green) and CifB (red) in the cytoplasmic waste bags (WB) that are present near the basal end of sperm tail bundles. Some of the Cif proteins strip down in the WB in *cifAB* line and absent in *w*Mel- control testes. Brightfield is shown to highlight the morphology of sperm tail bundles and waste bags, which are otherwise not visible using Cif antibodies and DAPI stain. The experiment was run in parallel to the ones shown in Fig 1 and Fig S2.

**Fig S6. Full uncropped fluorescent images are shown related to Fig 2A.**

**Fig S7. Individual CifA- and CifB-expressing lines do not show abnormal histone retention but are protamine deficient.** (A) Testes (n=15) from <8 hrs old males of single transgene-expressing lines *cifA*, *cifB* and a non CI-causing control gene *WD0508* were dissected to quantify spermatid bundles with histone retention (purple) during late canoe stage of spermatogenesis. DAPI stain (blue) was used to label spermatid nuclei. Total spermatid bundles with DAPI signals and those with retained Histones were manually counted and graphed. Single transgenic expressing lines showed significantly less histones similar to the negative control *w*Mel- at the late canoe stage. Vertical bars represent mean and error bars represent standard deviation. Letters indicate statistically significant (p<0.05) differences as determined by multiple comparisons based on a Kruskal-Wallis test and Dunn’s multiple test correction. (B) Mature sperms isolated from seminal vesicles (n=15) of <8 hr old males reared at 21°C were stained with fluorescent CMA3 (green) for detection of protamine deficiency in each individual sperm nucleus. Individual sperm head intensity was quantified in ImageJ (see methods) and graphed. *cifA*- and *cifB*-expressing lines showed significantly higher fluorescence indicative of reduced levels of protamines compared to *w*Mel- and *WD0508* control lines. Vertical bars represent mean and error bars represent standard deviation. Letters indicate statistically significant (p<0.05) differences as determined by multiple comparisons based on a Kruskal-Wallis test and Dunn’s multiple test correction. All of the P-values are reported in Table S1. The experiments were performed in parallel to the ones shown in Fig 2. (C) CI hatch rate analyses of transgenic male siblings used in CMA3 assays (Figure 2B, S6) validate that CI crosses (black circles) yielded significant less embryonic hatching compared to non CI-inducing ones, when reared at 21^0^C. Letters to the right indicate statistically significant (p<0.05) differences as determined by multiple comparisons based on a Kruskal-Wallis test and Dunn’s multiple test correction. All of the P-values are reported in Table S2. Raw data underlying this figure can be found in S1 data file.

**Fig S8. 7 day-old males do not cause CI and are not protamine deficient.** (A) Sperms from the 7 day-old wild type *Wolbachia*-infected (*w*Mel+) males show similar level of protamine levels as of *w*Mel-. Vertical bars represent mean, and error bars represent standard deviation. Letters indicate statistically significant (p<0.05) differences as determined by multiple comparisons based on a Kruskal-Wallis test and Dunn’s multiple test correction. All of the P-values are reported in Table S1. (B) CI hatch rate analyses of male siblings used in CMA3 assays (panel A) validate that 7d old *w*Mel+ do not induce CI that correlates with their normal levels of sperm protamine levels. Letters to the right indicate statistically significant (p<0.05) differences as determined by pairwise Mann-Whitney test and multiple comparisons calculated using a Kruskal-Wallis test and Dunn’s multiple test correction. All of the P-values are reported in Table S2. Raw data underlying this figure can be found in S1 data file.

**Fig S9. CifA is absent in late oocyte stages and the developing embryos from the rescue cross.** (A) In the transgenic *cifA* line, CifA (green) is absent in the late oocyte stages. Image was acquired at 20x magnification to show mid and late oocytes in one plane. We note the autofluorescence upon enhanced exposure in the green channel outlining the tissue morphology of stage 15 egg chamber does not signify CifA signals. (B) Immunofluorescence of CifA (green) and histones (magenta) in 1-2 hr old embryos (n=50) obtained from rescue cross (*cifAB* male x *w*Mel+ female). Histone signals are detected in the developing embryos colocalizing with host DNA, labelled with DAPI (blue), whereas CifA signals are absent.

