## Supplementary figures and images for "The Cif proteins from *Wolbachia* prophage WO modify sperm genome integrity to establish cytoplasmic incompatibility"

### Fig. S1.pdf

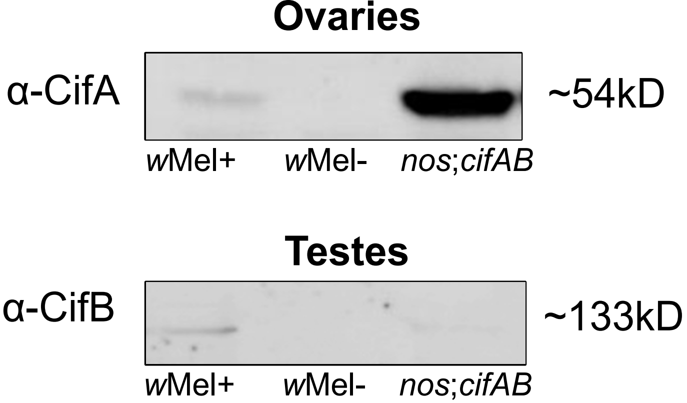


Uncropped images

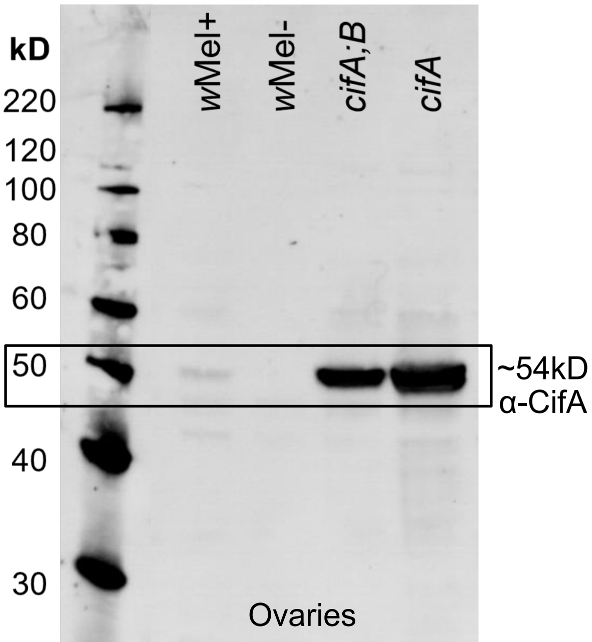


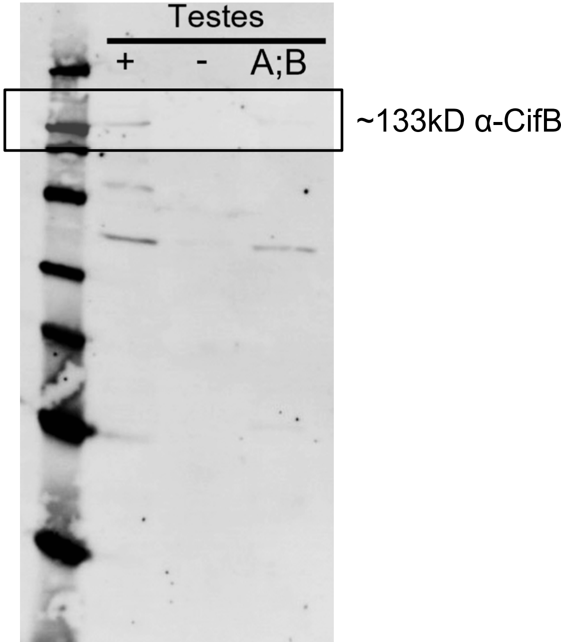

### Fig. S2.pdf

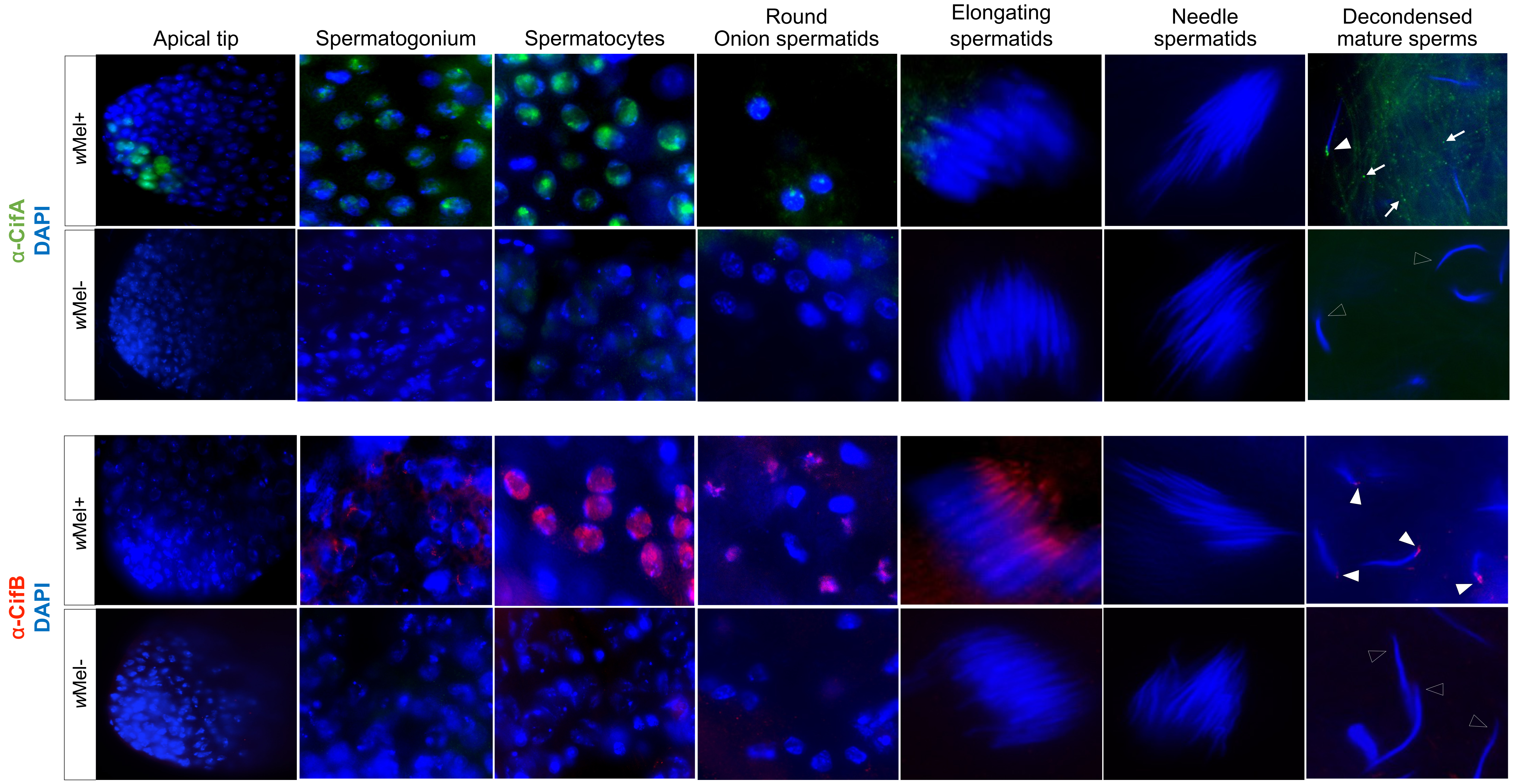

### Fig. S3.pdf

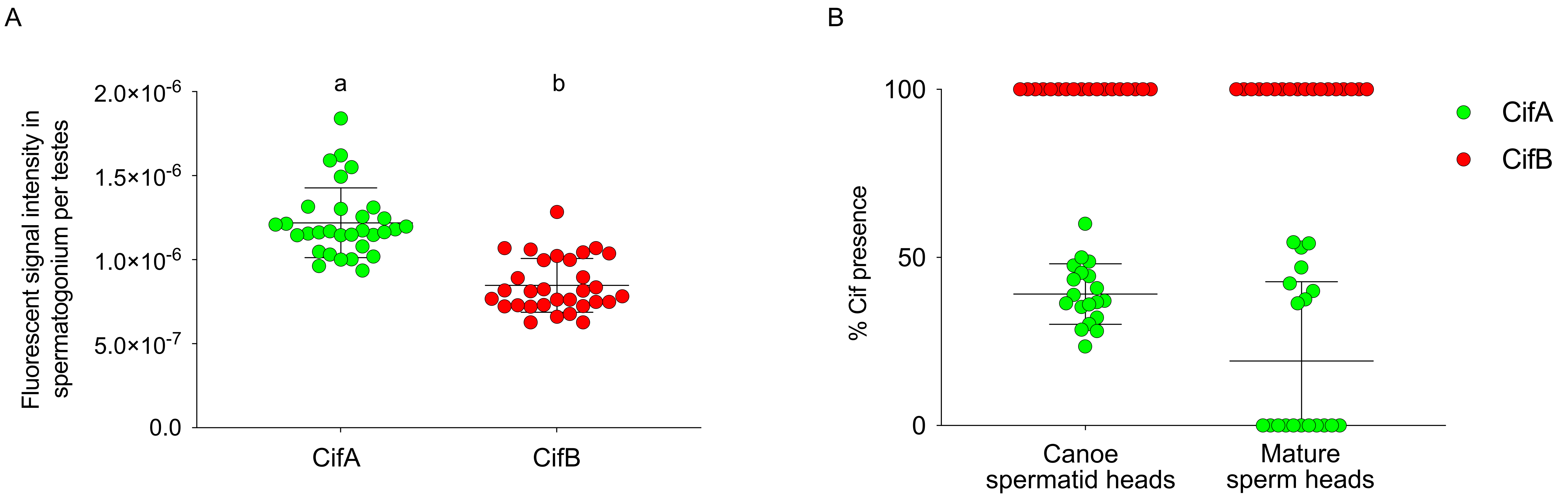

### Fig. S4.pdf

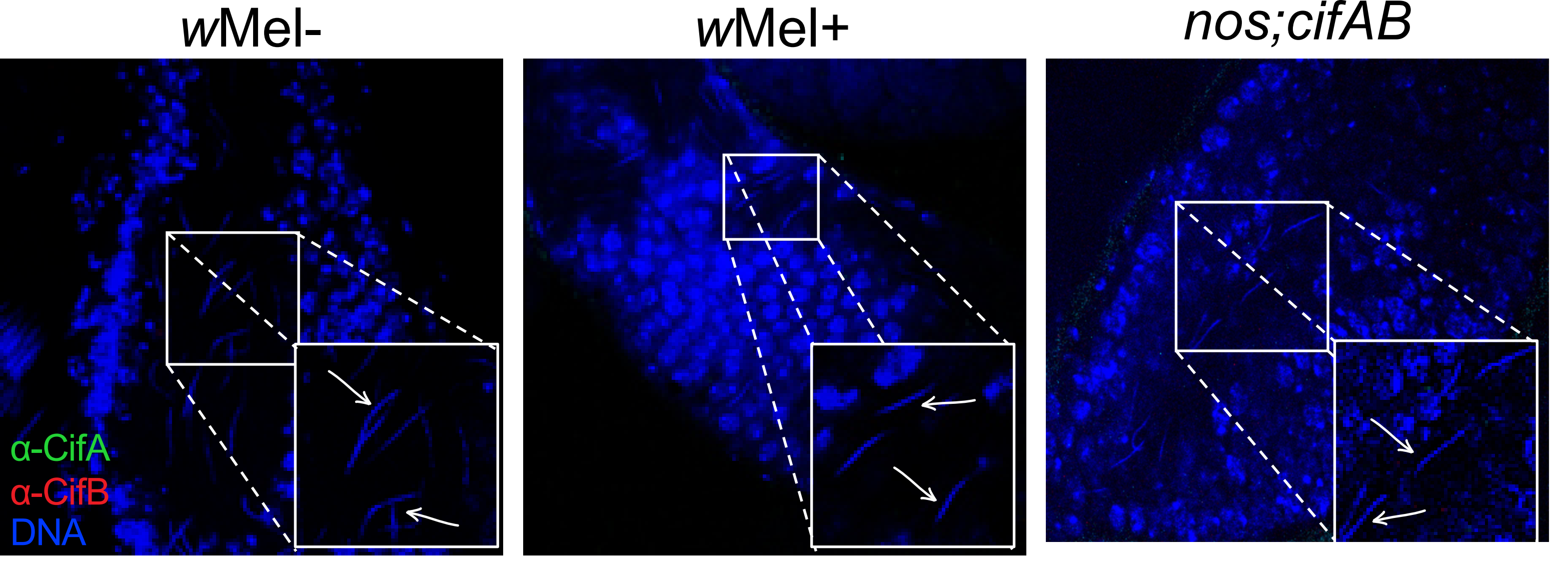

### Fig. S5.pdf

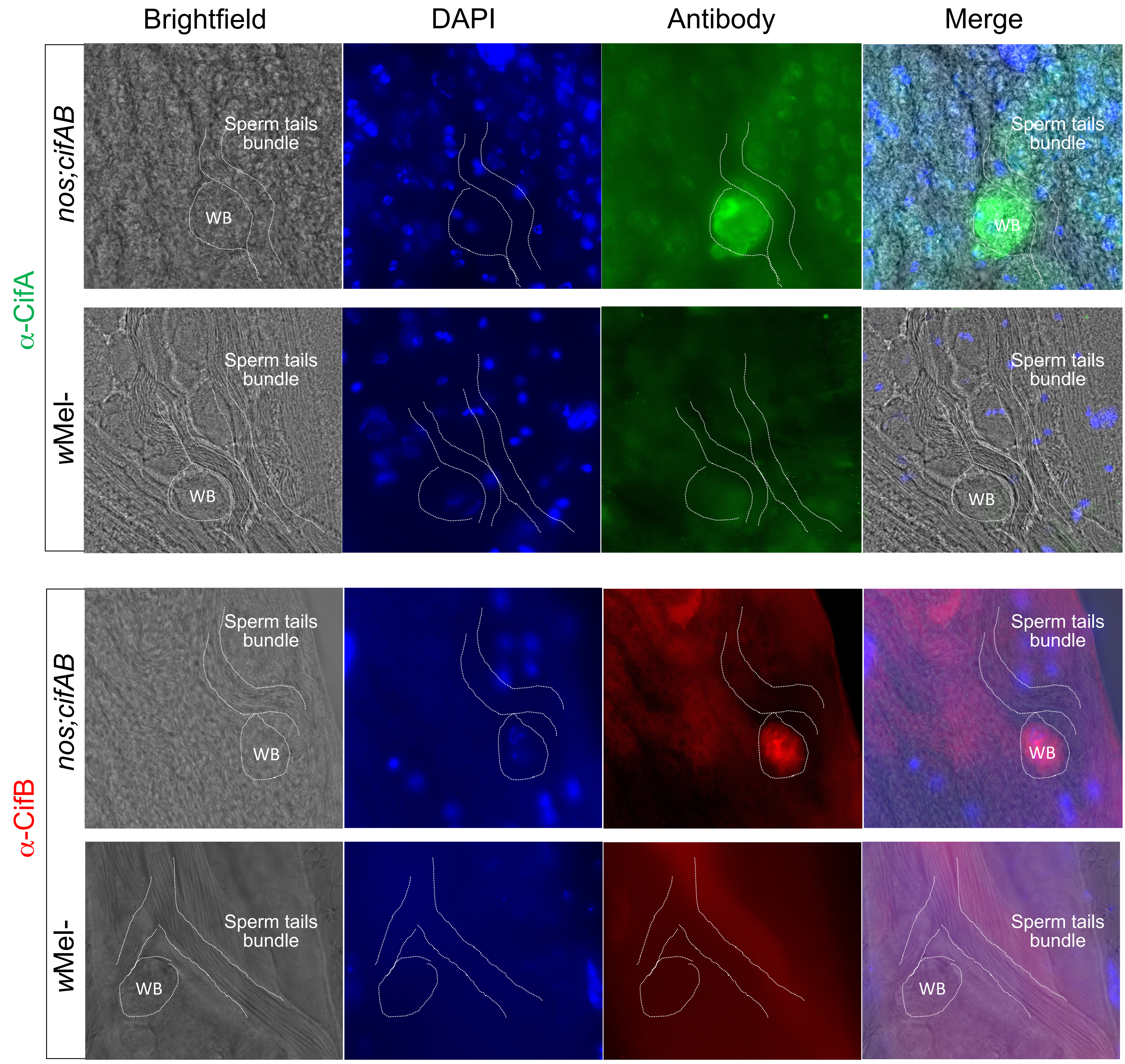

### Fig. S6.pdf

Uncropped images related to Fig. 2A

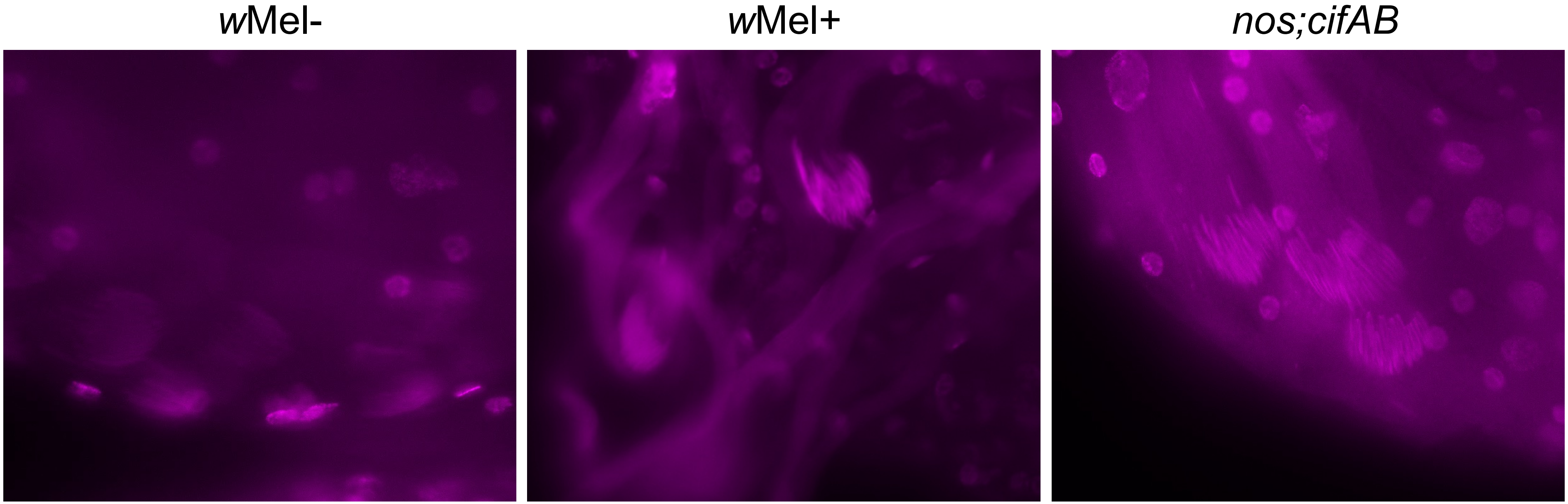

### Fig. S7.pdf

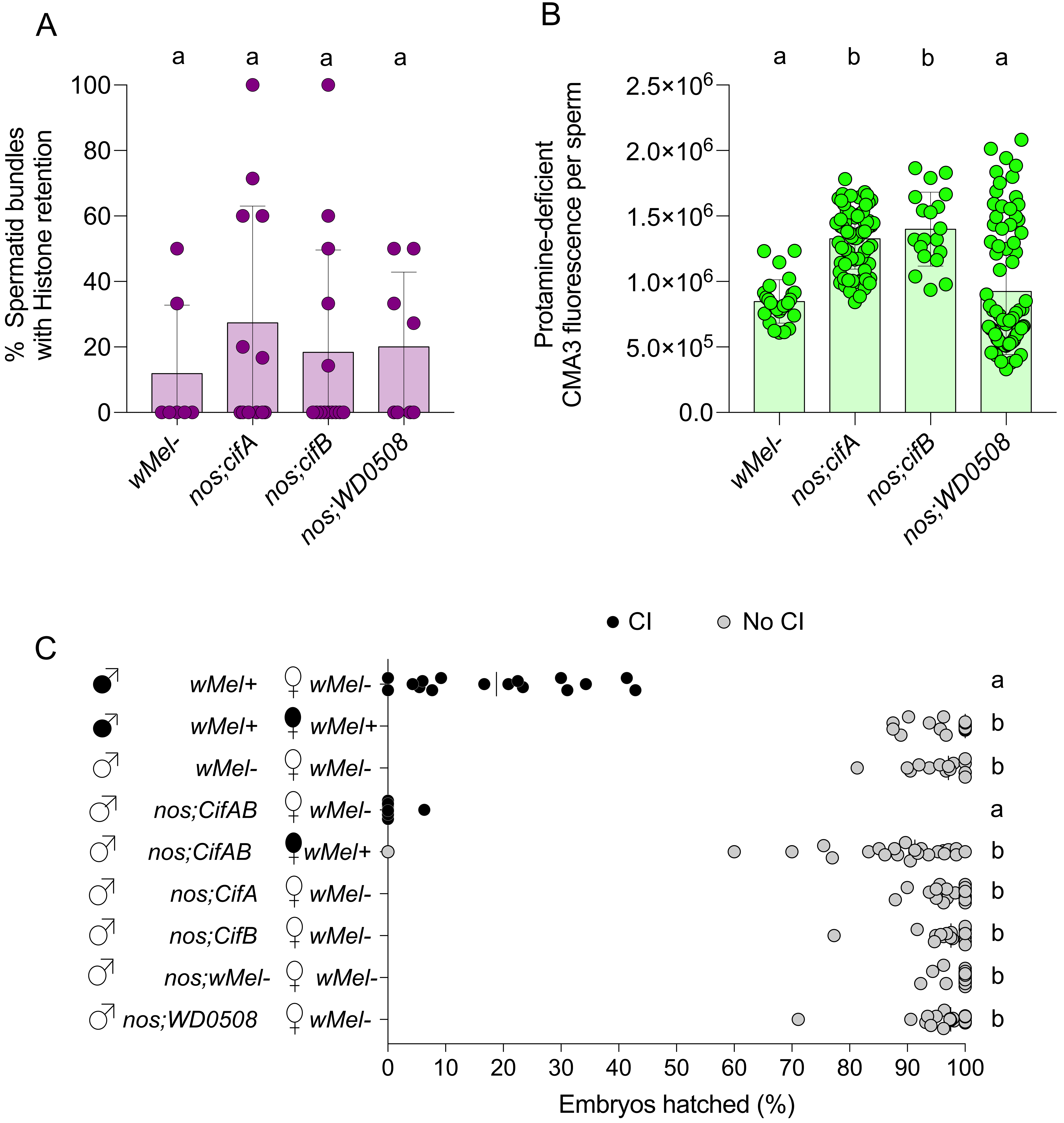

### Fig. S8.pdf

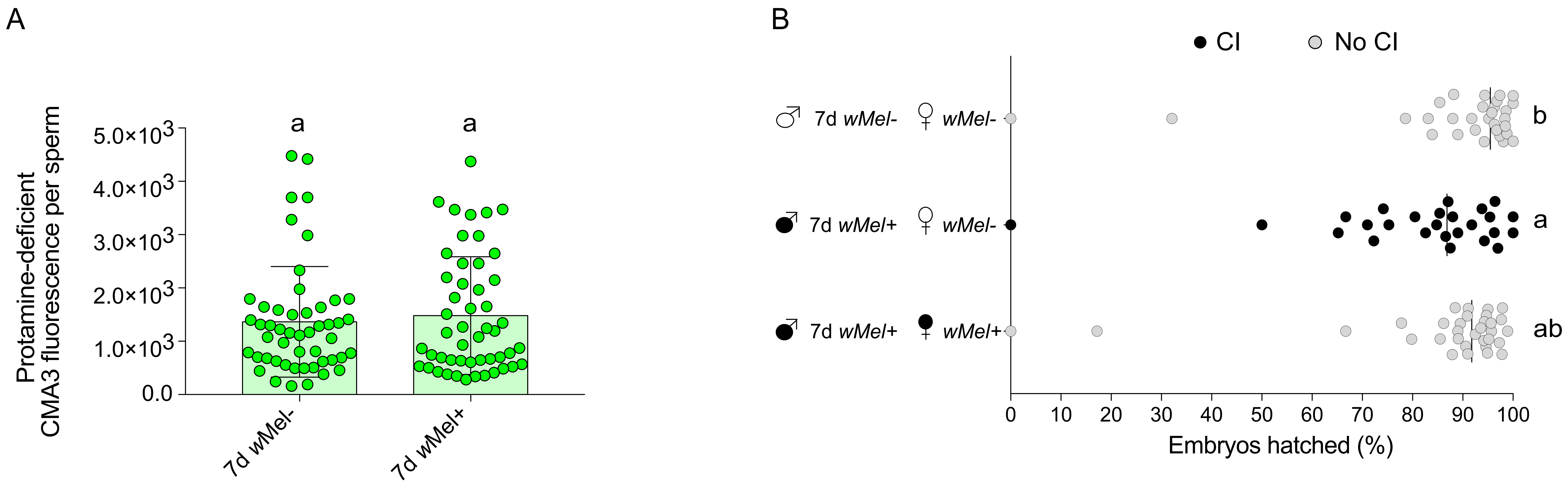

### Fig. S9.pdf

A

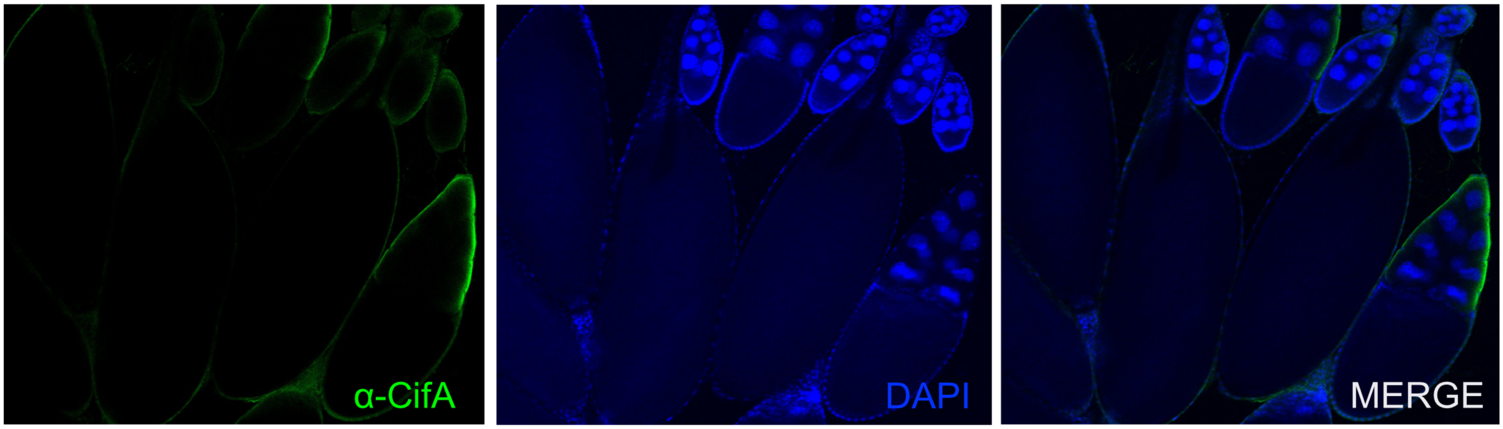


B

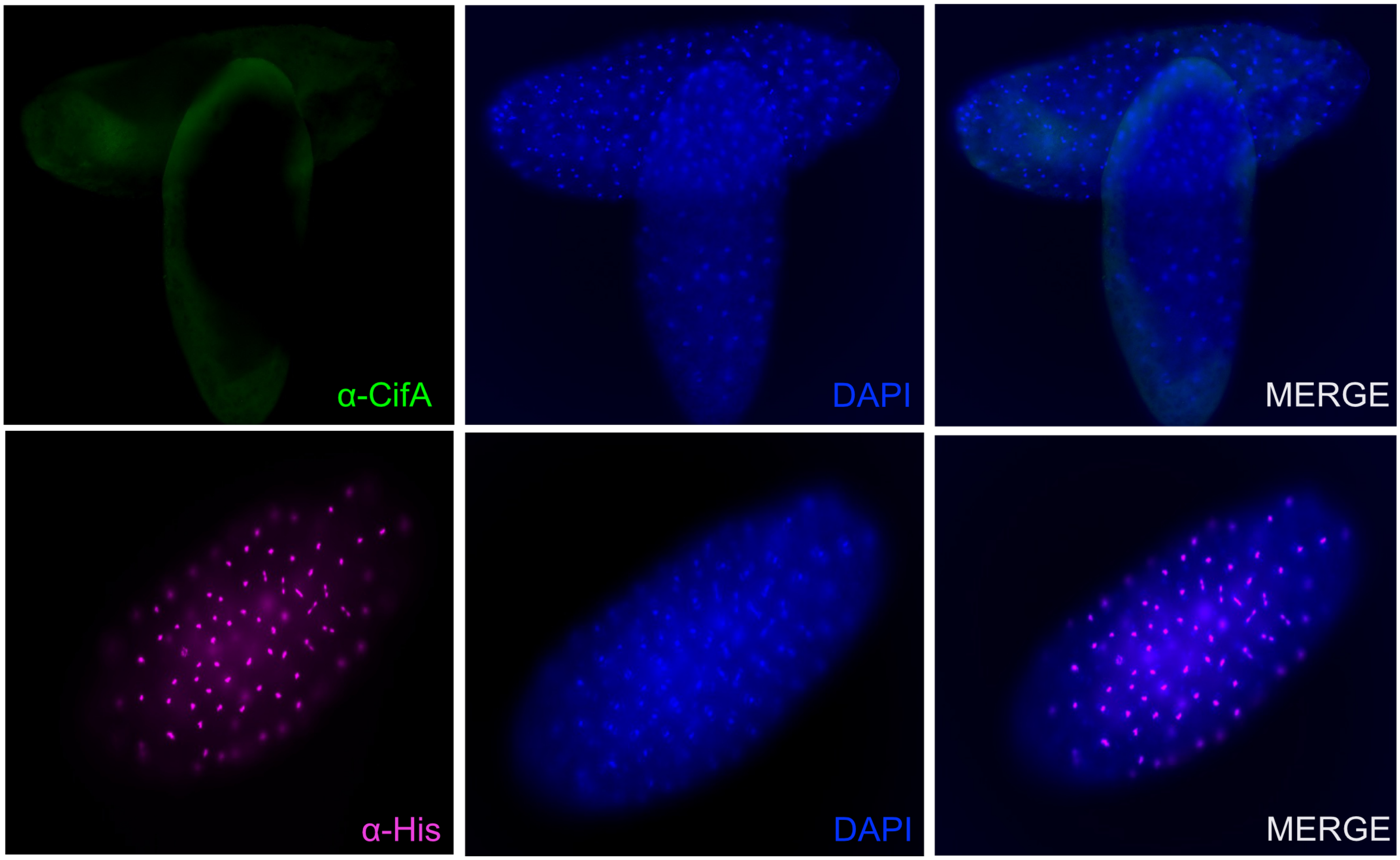
